# Molecular Evolution of Gustatory Receptors in the *Anopheles gambiae* Complex

**DOI:** 10.1101/2024.09.05.611554

**Authors:** Zachary R. Popkin-Hall, Michel A. Slotman

## Abstract

**Background:** Mosquitoes in the *Anopheles* (*An.*) *gambiae* species complex are major vectors of *Plasmodium falciparum* malaria. One reason for this is the high anthropophily of the constituent species *An. coluzzii*, *An. gambiae* sensu stricto, and *An. arabiensis*. In contrast, their sister species *An. quadriannulatus* is highly zoophilic. *Anopheles* mosquitoes largely rely on chemical cues for host-seeking, which are primarily detected by four chemosensory gene families: olfactory receptors (*Ors*), ionotropic receptors (*Irs*), gustatory receptors (*Grs*), and odorant binding proteins (*Obps*). Genes from these families that have been implicated in host adaptation show evidence of positive selection in other insect species, including other mosquitoes. As such, we analyzed the molecular evolutionary patterns of the gustatory receptors within the *Anopheles gambiae* complex, with a particular interest in identifying *Grs* that show evidence of positive selection in highly anthropophilic species.

**Results:** We identified sixteen *Grs* that show evidence of potential positive selection using the McDonald-Kreitman test, including four putative sugar receptors and two *Grs* with unknown ligands that are relatively highly expressed in chemosensory organs of either *An. coluzzii* or *An. quadriannulatus*. In addition, we identified twelve *Grs* that show evidence of potential purifying selection using the McDonald-Kreitman test, and twelve *Grs* that may have experienced a selective sweep using the DH test, including three putative sugar receptors and the carbon dioxide receptor *Gr24*. We also identified both positive and purifying selection in the coastal species *An. melas* (West Africa) and *An. merus* (East Africa).

**Conclusions:** Our results, together with transcriptomic data, identify four *Grs* as possible candidates for involvement in the evolution of vertebrate host preference in the *An. gambiae* complex, as may have occurred in the *An. farauti* complex. They also point to sugar receptors as playing a role in recent adaptation of some of these species. As the vast majority of *Grs* have unknown functions and much is still unknown about the role of *Grs* in these species, a more complete interpretation of our data necessitates further characterization of these genes.

## Background

Mosquitoes in the *Anopheles* (*An.*) *gambiae* complex are the major vectors of malaria in sub-Saharan Africa, and consequently are responsible for the deaths of hundreds of thousands of children every year. One reason for this high mortality is the high anthropophily of two complex members (*An. coluzzii* and *An. gambiae* s.s.). As such, elucidating the genetic basis of their human host preference is important for understanding a critical aspect of the biology of these species, but may also provide candidate genes for developing novel control methods. Because chemosensory perception plays a crucial role in host seeking and preference, these gene families have been studied extensively in mosquitoes, particularly through RNA-seq studies [1–12] and their ligand binding properties [13–21].

In insects the chemosensory genes comprise four major gene families: the gustatory receptors (*Grs*), ionotropic receptors (*Irs*), odorant binding proteins (*Obps*), and olfactory receptors (*Ors*), in addition to several others that play minor roles [22–26]. The GRs, ORs and IRs form ligand activated ion channels [27, 28]. Unlike ORs, which form heterodimers of the obligate olfactory co-receptor (ORCO) and one specific OR [29], GRs can be multimeric and tend to be co-expressed with multiple other GRs [30], with two to four gustatory neurons per gustatory sensillum [31, 32] and two to five GRs per gustatory neuron [33]. GRs mediate perception of a wide variety of chemical cues, the majority of which are concentrated tastants, rather than airborne volatile chemicals [34].

However, three *Grs, AgGrs22-24,* together encode the carbon dioxide receptor in An. coluzzii [35]. These receptors are highly conserved among the major disease vectors. Recent work on *Aedes* CO_2_ receptors suggests that only two of these receptors are necessary to detect CO_2_, while the third (the *AgGr22* homolog *AeGr1*) also detects other molecules [36].Other GRs respond to sugars, salts, pheromones, and bitter compounds in complex ways, with different combinations of *Grs* eliciting specific responses to particular concentrations of chemicals, and sometimes inhibiting one another [31, 32, 37–42].

Studies in *Drosophila* have localized *Gr* expression to the taste organs, the brain, olfactory neurons, digestive tract, as well as non-chemosensory neurons, providing evidence of non-gustatory roles [40, 43–51]. Although many *Grs* have been deorphanized in *Drosophila*, the vast majority of anopheline *Grs* have unknown ligands. The functional characterization of *Grs* has been more difficult than for other receptors, due to the inability to heterologously express these genes [52]. As such, while the carbon dioxide receptors and some sugar receptors have been identified in *Anopheles*, the biological function of most anopheline *Grs* remains unclear.

*Grs* are an ancient gene family, being already present in such basal animals as placozoans [53, 54]. The *Grs* are therefore the basal member of a larger insect chemosensory receptor superfamily that also includes the *Ors*, which are derived from within the *Grs* [55, 56]. This superfamily is characterized by large numbers of lineage-specific expansions throughout Arthropoda [53, 57, 58]. The *Gr* repertoire tends to be smallest in specialists and largest in generalists [59], and can vary substantially within orders and even genera. Although there are numerous pseudogenizations documented in *Drosophila* (reviewed in Robertson (2019)), the *Gr* repertoires of both *Anopheles* and another hematophagous fly genus, *Glossina*, are much more stable [60, 61].

The evolution of the insect chemosensory gene superfamily tends to follow a birth-and-death model with numerous lineage-specific duplications, followed by pseudogenization and eventual gene loss [57, 62]. While *Ors* are derived from *Grs*, the evolutionary dynamics of these two gene families are not identical: *Grs* have increased replacement divergence (i.e. fixed amino acid changes between species) relative to *Ors*, as well as lower neutrality indices. This difference could stem from either stronger positive selection or weaker purifying selection in *Grs* vs *Ors* [63, 64]. *Grs* likely underwent gene duplications followed by subsequent differentiation following speciation events, and exhibit low sequence similarities to one another both within and between species [53, 55].

Positive selection during host shifts associated with speciation has been detected in the *Grs* of several insect taxa, including multiple *Drosophila spp.* [63, 65, 66], the butterfly *Heliconius melpomene* [67], the pea aphid *Acyrthosiphon pisum* [68], and may have combined with gene family expansion to facilitate adaptive radiation throughout Lepidoptera [69]. For example, in the butterfly *Vanessa cardui*, frequent *Gr* duplications occurred in the transition from a specialist to a generalist lifestyle [70]. *Gr* plasticity in responding to environmental change and a role of mutations in *Grs* in modulating species-specific behaviors is also found in *Drosophila spp.* [71], the German cockroach *Blattella germanica* [72], and the butterfly *Papilio xuthus* [73].

Nonetheless, positive selection on the whole is thought to play a relatively minor role in the evolution of *Grs*, with strong purifying selection acting as the dominant evolutionary force in *Drosophila* and Lepidoptera [57, 74, 75] while most diversification, particularly in the *Grs*, appears to be due to relaxed purifying selection, rather than positive selection [64]. In addition, selection on chemosensory gene expression may be responsible for altering sensitivity to certain odors, rather than changes in the protein structure resulting in changes in ligand binding affinity [76]. Numerous chemosensory genes, including some *Grs*, are differentially expressed between members of the *An. gambiae* complex with different host preferences [2, 9, 12], which could reflect differential sensitivity to odors. Intriguingly, work in *Aedes aegypti* has demonstrated that differential expression as well as nucleotide substitutions in a single *Or* can influence host preference between two closely related subspecies [77]. In addition, 22 genes, including six *Grs*, show evidence of involvement in vertebrate host preference in the *An. farauti* complex [78]. One of these *Grs* is a strong candidate for anthropophily[78].

The *An. gambiae* complex consists of nine cryptic species with varying host preferences and distributions, six of which are included in this study: the highly anthropophilic major vectors *An. coluzzii* (formerly M form) and *An. gambiae* sensu stricto (formerly S form) [79–86]; *An. arabiensis*, which is also anthropophilic and a major vector, but which exhibits substantial host choice plasticity and prefers dryer habitats [85, 87–90]; the range-limited locally important vectors *An. melas* and *An. merus*, which are found in brackish habitats in West and East Africa, respectively, and are less anthropophilic than the others [90–97]; and finally the non-vector *An. quadriannulatus*, which feeds primarily on non-human animals, particularly cattle [82, 98–101]. Because *An. arabiensis* is both a major vector and a more generalist feeder, multiple studies have assessed this species for genes that could be involved in human host preference, and have identified genes within inversions on chromosomes 2R and 3R that are associated with differences in vertebrate host preference [102–104]. There is also substantial introgression primarily of autosomal regions between *An. arabiensis*, *An. coluzzii*, and *An. gambiae* s.s. [105, 106], as well as between *An. merus* and *An. quadriannulatus* [107]. Selection maintains the 2La inversion in *An. arabiensis* [108, 109], and it is likely that the other genes which have introgressed into *An. arabiensis* are adaptive for its shift to anthropophily [107].

Previous work has focused on differential expression of chemosensory genes in the major mosquito sensory organs between the anthropophilic *An. coluzzii* and zoophilic *An. quadriannulatus* [2, 9, 12], as well as the physical ablation of said organs [110–115]. In the present study, we survey the molecular evolutionary patterns of the *Grs* in the *An. gambiae* complex in the context of species-specific host-seeking behaviors. We extracted whole *Gr* sequences from each of our six focal species from the 16 *Anopheles* genomes project [61], and screened for evidence of selection using the McDonald-Kreitman test [116] and direction of selection (DoS)[117], as well as the DH and E tests [118]. Most *Grs* in the *An. gambiae* complex have unknown ligands, with the exception of the CO_2_ receptors listed above, the putative sugar receptors *Grs* 14-21 [119], and the sugar receptor *Gr25* [120]. We are particularly interested in identifying *Grs* that diverged between the anthropophilic *An. coluzzii*/*An. gambiae* s.s. clade and the zoophilic *An. quadriannulatus*. Furthermore, we describe signatures of positive selection in other lineages, as well as evidence of purifying selection and recovery from selective sweeps. We focus especially on genes with notable expression levels in the chemosensory organs of *An. coluzzii* and *An. quadriannulatus*, We identify potential evidence of selection in four putative sugar receptors, two *Grs* with unknown functions that are highly expressed in chemosensory organs of male *An. quadriannulatus*, as well as several *Grs* with unknown functions and unknown expression patterns. These results point to multiple avenues for further detailed exploration of anopheline *Grs*.

## Methods

### Data

Whole genome sequences for six members of the *An. gambiae* complex (*An. arabiensis*, *An. coluzzii*, *An. gambiae* s.s., *An. melas*, *An. merus*, and *An. quadriannulatus*) from the 16 *Anopheles* Genomes Project [61, 121] were downloaded from NCBI. No comparable data exist for the remaining constituent species of the complex, none of which has been extensively studied. Genome data for a total of 96 *An. arabiensis*, 12 *An. coluzzii*, 26 *An. gambiae*, 65 *An. melas*, 72 *An. merus*, and 72 *An. quadriannulatus* specimens were used for analysis.

### Variant Discovery Pipeline

Whole genome sequences were processed according to the GATK 4 Best Practices Workflow for DNA Variant Discovery [122]. Reads were aligned to their respective genomes in BWA [123], with the exception of *An. coluzzii*, which was aligned to the *An. gambiae* genome due to the lower quality of the *An. coluzzii* genome and the their mostly shared genetic make-up. All genomes were downloaded from VectorBase [124].

Following mapping in BWA, Picard Tools (http://broadinstitute.github.io/picard) was used to add read group information and mark duplicate reads. A bed file was created to mark intervals corresponding to the genomic coordinates of the genes of interest, which were determined by running a local BLAST search [125] using the *An. gambiae* gene sequences. In cases where more than one nonconsecutive match was recovered, regions were prioritized by length, e-value, and bit score.

This interval file was used to restrict the use of HaplotypeCaller to only the genes of interest, thereby increasing processing speed. Haplotype Caller was run using gVCF mode to improve the accuracy of physical phasing where possible, and all samples from a given species were processed together through the remaining steps. Indels were excluded from further analysis, as were any reads not meeting the following quality standards: QD < 2.0 || FS >55.0 || MQ < 40.0 || MQRankSum < −12.5 || ReadPosRankSum < −8.0.

A custom bash script incorporating bcftools [126] was used to screen for low coverage sequences by counting the number of variants genotyped in the entire species dataset, and then comparing individual sequences for the presence of those genotypes (whether variant or non-variant). All individuals with an average of at least 50% of variants successfully genotyped across all genes were used for McDonald-Kreitman analysis, while sequences with missing genotypes were excluded from DH test analyses. bcftools was also used to extract two phased fasta sequences for each individual. BEDTools [127] was used to extract individual genes from each genome sequence. Two data sets were generated using the masked gene data sets: one with full gene sequences including introns and another including only coding sequence (CDS). In the case of genes with multiple splice variants, multiple CDS data sets were generated.

### Data Analysis

Sequences were obtained for all species for *AgGrs1-60* with the following exceptions: a complete *Gr6* sequence was not found in the *An. arabiensis* genome; complete *Gr2, Gr10*, *Gr44*, *Gr56*, and *Gr57* sequences were not found in the *An. Melas* genome; complete *Gr44*, *Gr49*, *Gr53* and *Gr57* sequences were not found in the *An. merus* genome. *Gr5*, *Gr9*, and *Gr10* were not analyzed due to a low number of available sequences and the presence of premature stop codons.

CDS data sets including all individuals were imported into DnaSP version 6.12.03 [128], which was then used to perform a McDonald-Kreitman test [116] between each pair of species to detect signs of positive selection. All sequences were used for this test, as it relies solely on comparisons between polymorphic sites in coding sequences. Different splice variants were treated as unique genes for this test, but not for any others which do not rely on coding sequences only. *Grs* were considered to be undergoing positive selection if a significant p-value was calculated in addition to a direction of selection (DoS) value > 0.01, whereas they were considered to be undergoing purifying selection if a significant p-value was combined with DoS < −0.01.

*Grs* with DoS between −0.01 and 0.01 were considered to be undergoing neutral evolution. These conservative cutoffs were chosen based on simulated performance of DoS[117]. DoS is less susceptible to bias than the neutrality index, particularly when estimates are based on relatively few SNPs.DoSt is calculated as, DoS = *D*_n_/(*D*_n_ + *D*_s_) − *P*_n_/(*P*_n_ + *P*_s_) where *D*_n_ refers to the number of fixed replacement substitutions between species, *D*_s_ refers to the number of fixed synonymous substitutions between species, *P*_n_ refers to the number of polymorphic replacement substitutions within a species, and *P*_s_ refers to the number of polymorphic synonymous substitutions within a species. DnaSP was also used to generate basic population parameters for each species by the use of a Fu and Li D test [129], from which only the population parameters and not the significance values were considered.

Datasets including the full gene sequence were produced only if at least ten sequences with known genotypes for each variant site existed in a given species. In cases of alternatively spliced genes, the longest possible sequence was used for the DH test. As a result of these more stringent criteria, these data sets were primarily produced for *An. coluzzii* and *An. gambiae* s.s., which were sequenced at a higher depth of coverage than the other species in this study. These data sets were exported in fasta format with one outgroup sequence. Where possible, data sets were produced with *An. coluzzii* and *An. gambiae* s.s. analyzed both as individual species and as a single clade. These fasta files were loaded in the *Readms* module of DH (available from https://github.com/drkaizeng/publications-and-software/blob/main/dh/dh.zip), where the following tests were implemented with 10,000 coalescent simulations: D [130], normalized H [118, 131], DH [118], and E [118].

While all of these tests can detect directional selection, DH is unique in its insensitivity to other population genetic forces and is designed to detect evidence of selective sweeps. Tajima’s D can detect balancing or purifying selection but is also sensitive to changes in population size. Fay and Wu’s H is used primarily to detect genetic hitchhiking but is also sensitive to reductions in population size and the presence of population structure. The E test identifies the recovery of genetic diversity following its loss (e.g. following a selective sweep) and is robust to population structure, but sensitive to both background selection and increases in population size [118].Finally, TCS haplotype networks were generated in POPART[132].

## Results

A total of 79 to 260 sequences were obtained for most of the *AgGrs1-60* for the six species (**Supplemental Table 1**). All *Grs* for which data was available were included in our analyses and data tables, but only those *Grs* which fulfill the following criteria are discussed in the text: a significant or nearly significant test result in one of our focal species (*An. coluzzii*, *An. gambiae*, *An. quadriannulatus*, and to a lesser extent, *An. arabiensis*), and either a) a known or suspected function, or b) high or differential expression in the chemosensory organs of *An. coluzzii* and/or *An. quadriannulatus*.

### Fixed Differences Between Species

No fixed differences were identified between *An. coluzzii* and *An. gambiae* in any *Gr* coding sequence (**Table 1**). Similarly, no fixed differences were found in the majority of *Grs* between either of these species and *An. arabiensis* (53 out of 75 *Grs* for *An. coluzzii*, and 60 out of 75 *Grs* for *An. gambiae*). However, the majority of *Grs* (between 64 and 71) have fixed differences between these three species and *An. quadriannulatus*. Furthermore, fixed differences are present at almost every *Gr* locus between every species pair that includes either *An. melas* or *An. merus*.

**Table 1.** Grs with fixed differences (both synonymous and non-synonymous) between species pairs.

| Species Pair | # of Grs with Fixed Differences | # of Grs without Fixed Differences | # of Grs Analyzed | Percent Grs with Fixed Differences |
| --- | --- | --- | --- | --- |
| *arabiensis-coluzzii* | 22 | 53 | 75 | 29.3% |
| *arabiensis-gambiae* | 15 | 60 | 75 | 20.0% |
| *arabiensis-melas* | 62 | 0 | 62 | 100% |
| *arabiensis-merus* | 66 | 1 | 67 | 98.5% |
| *arabiensis-quadriannulatus* | 71 | 4 | 75 | 94.7% |
| *coluzzii-gambiae* | 0 | 76 | 76 | 0% |
| *coluzzii-melas* | 63 | 0 | 63 | 100% |
| *coluzzii-merus* | 67 | 1 | 68 | 98.5% |
| *coluzzii-quadriannulatus* | 64 | 12 | 76 | 84.2% |
| *gambiae-melas* | 63 | 0 | 63 | 100% |
| *gambiae-merus* | 67 | 1 | 68 | 98.5% |
| *gambiae-quadriannulatus* | 65 | 11 | 76 | 85.5% |
| *melas-merus* | 60 | 0 | 60 | 100% |
| *melas-quadriannulatus* | 63 | 0 | 63 | 100% |
| *merus-quadriannulatus* | 67 | 0 | 67 | 100% |

### McDonald-Kreitman Test

Next, the *Grs* were analyzed using the McDonald-Kreitman (MK) test between every species pair (**Supplemental Table 2**). Because the large number of tests conducted in this study precludes the conclusive identification of positive selection due to the multiple testing problem, all significant MK results are suggestive, but not definitive evidence of selection. While the Benjamini-Hochberg procedure can be used to produce an adjusted p-value and reduce type I error rate, it also substantially decreases power. This is particularly problematic in closely related species pairs with low numbers of fixed differences, in which the power of the MK test is low to begin with. Therefore, unadjusted p-values are given here with the caveat noted above.

When comparing the anthropophilic *An. arabiensis*, *An. coluzzii*, and *An. gambiae* s.s. to the zoophilic *An. quadriannulatus*, most *Grs* show signatures of purifying selection (49.3%, 60.9%, and 55.4%, respectively) (**Figure 1A**, **Table 2**). By comparison, signatures of positive selection are detected in 45.5%, 37.5%, and 43.1% of *Grs*, respectively (**Figure 1A**, **Table 2**). As indicated in **Table 1**, there are few fixed differences between *An. arabiensis* and either *An. coluzzii* or *An. gambiae* s.s. In both comparisons, the *Grs* with fixed differences are approximately evenly split between showing signatures of positive selection and signatures of purifying selection (**Table 2**, **Figure 1B**).

**Figure 1.**
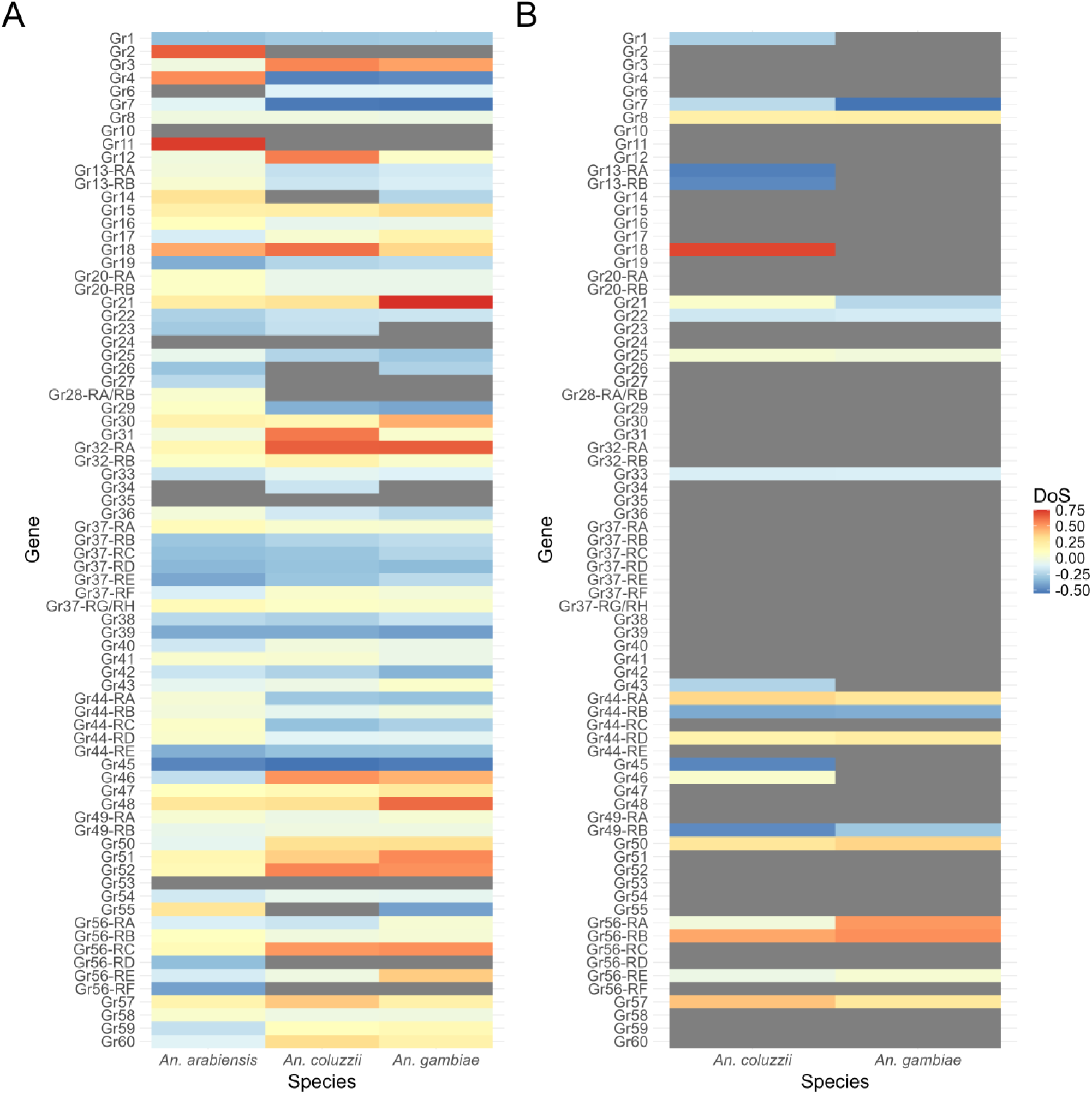
Heat map of direction of selection (DoS) between **A)** the three major vector species vs. An. quadriannulatus and **B)** An. coluzzii and An. gambiae s.s vs. An. arabiensis. Positive values (consistent with positive selection) are red, while negative values (consistent with purifying selection) are blue and neutral values are yellow. Grs where NI could not be computed because of a lack of fixed non-synonymous differences are represented in grey.

**Table 2.** Prevalence of positive and purifying selection based on Direction of Selection between species pairs. Genes with no fixed differences are excluded from these counts.

| **<u>Species Pair</u>** | **<u>Grs with DoS > 0.01</u>**<br>**<u>(Positive Selection)</u>** | **<u>Grs with DoS ≥ -0.01</u>**<br>**<u>& ≤ 0.01</u>**<br>**<u>(Neutral Evolution)</u>** | **<u>Grs with NI < -0.01</u>**<br>**<u>(Purifying Selection)</u>** |
| --- | --- | --- | --- |
| *arabiensis-quadriannulatus* | 32 | 4 | 35 |
| *coluzzii-quadriannulatus* | 24 | 1 | 39 |
| *gambiae-quadriannulatus* | 28 | 1 | 36 |
| *arabiensis-coluzzii* | 10 | 0 | 12 |
| *arabiensis-gambiae* | 8 | 0 | 7 |
| *arabiensis-melas* | 18 | 9 | 35 |
| *arabiensis-merus* | 30 | 1 | 35 |
| *coluzzii-melas* | 32 | 4 | 27 |
| *coluzzii-merus* | 24 | 2 | 41 |
| *gambiae-melas* | 35 | 3 | 25 |
| *gambiae-merus* | 24 | 4 | 39 |
| *melas-merus* | 11 | 4 | 45 |
| *melas-quadriannulatus* | 27 | 5 | 31 |
| *merus-quadriannulatus* | 28 | 2 | 37 |

When comparing *An. melas to An. merus*, the majority of DoS values are negative, i.e. consistent with purifying selection (75.0%) (**Table 2**, **Figure 2A**). When comparing *An. melas* to the three anthropophilic species, purifying selection is more common vs. *An. arabiensis* (56.5% vs. 29.0% with positive DoS), but positive selection is more common vs. both *An. coluzzii* (50.8% vs. 42.9% with negative DoS) and *An. gambiae* s.s. (55.6% vs. 39.7% with negative DoS) (**Table 2**, **Figure 2**). *An. merus* consistently shows a higher proportion of negative DoS values when compared to the three anthropophilic species (53.0% vs. *An. arabiensis*, 61.2% vs. *An. coluzzii*, 58.2% vs. *An. gambiae* s.s.) (**Table 2**, **Figure 2**). Finally, when comparing *An. melas* and *An. merus* to *An. quadriannulatus*, the most *Grs* again have negative DoS values consistent with purifying selection (49.2% and 55.2%, respectively) (**Table 2**, **Figure 2**).

**Figure 2.**
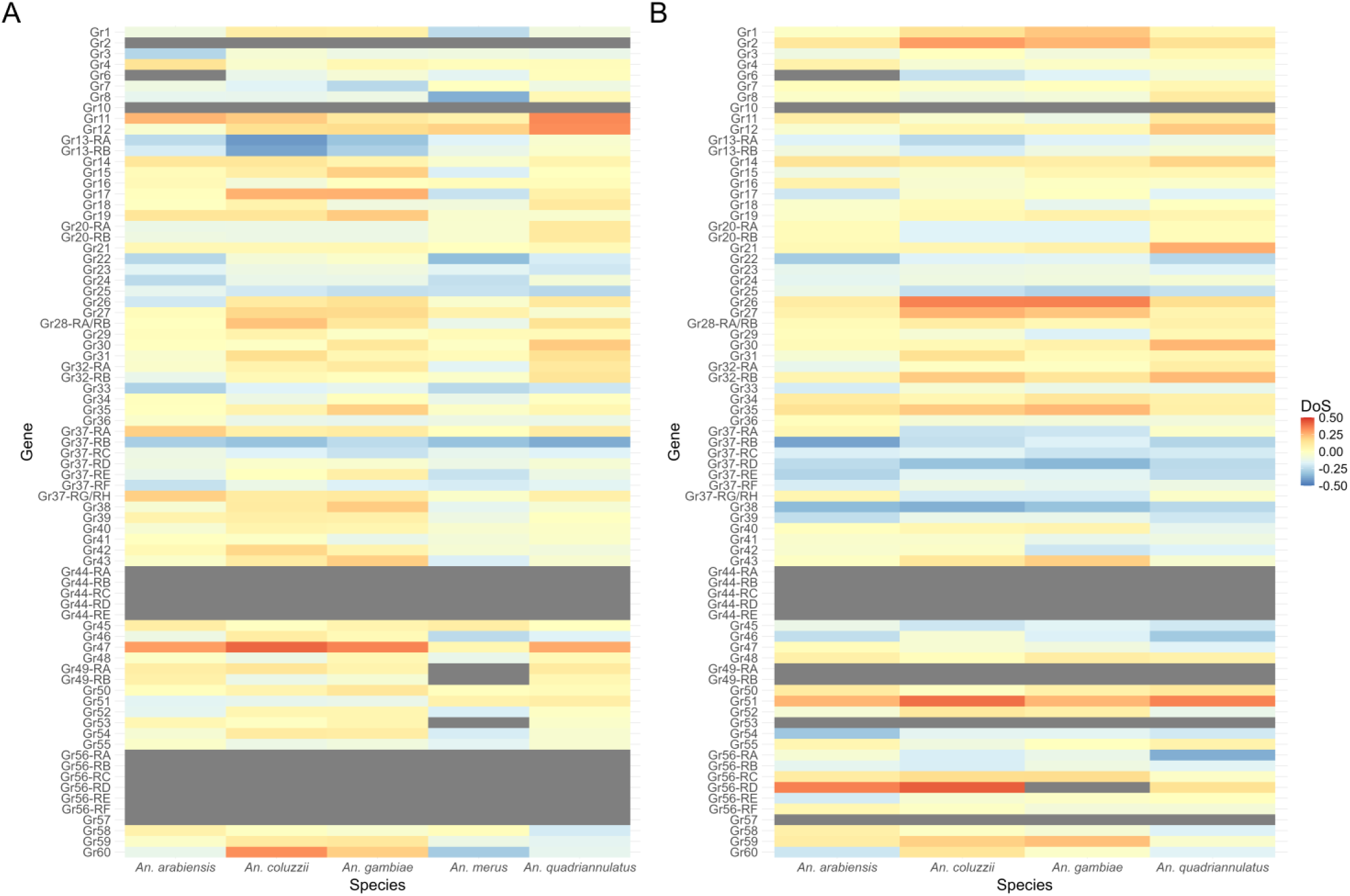
Heat map of neutrality indices (NI) between **A)** An. melas vs. the other complex species and **B)** An. merus vs. the other complex species. Positive values (consistent with positive selection) are red, while negative values (consistent with purifying selection) are blue and neutral values are yellow. Grs where NI could not be computed because of a lack of fixed non-synonymous differences are represented in grey.

Genes showing evidence of positive selection when comparing the anthropophilic *An. coluzzii* and *An. gambiae* s.s. and the zoophilic *An. quadriannulatus* are of particular interest as these are candidates to play a role in the divergent host preference between these species. We identify putative positive selection as an excess of fixed replacement substitutions according to a significant p-value (< 0.05) on the MK test, with the magnitude indicated by a positive DoS value. Only one of 76 *Gr*s was indicated to be under positive selection between these species pairs: the sugar receptor *Gr18* in the *An. coluzzii* – *An. quadriannulatus* comparison (**Table 3**). Three additional *Grs* had marginally significant (0.05 < p < 0.07) excesses of fixed replacement substitutions: *Gr21, Gr48*, *and Gr60* (**Table 3**)*. An. arabiensis* is also of interest as a relatively anthropophilic close relative of *An. quadriannulatus* with extensive signatures of introgression with *An. coluzzii* and *An. gambiae* s.s. [133, 134]. Three *Grs* showed excesses of fixed replacement substitutions between *An. arabiensis* and *An. quadriannulatus*: *Gr4*, *Gr18* (both significant, **Table 3**), and *Gr48* (marginally significant).

**Table 3.** Grs with significant (or near-significant) excesses of fixed substitutions and positive direction of selection (DoS), suggestive of positive selection.

| **Gene** | **Species Pair** | **dN** | **dS** | **dN/dS** | **pN** | **pS** | **pN/pS** | **p-value** | **DoS** |
| --- | --- | --- | --- | --- | --- | --- | --- | --- | --- |
| Gr2 | coluzzii-merus | 8 | 7 | 1.14 | 18 | 60 | 0.3 | 0.027 | 0.302 |
| Gr4 | arabensis-quadriannulatus | 5 | 0 | NA | 57 | 67 | 0.851 | 0.024 | 0.540 |
| Gr11 | arabensis-melas | 16 | 11 | 1.45 | 17 | 33 | 0.515 | 0.053 | 0.253 |
| Gr11 | melas-quadriannulatus | 15 | 12 | 1.25 | 9 | 33 | 0.273 | 0.005 | 0.341 |
| Gr12 | melas-quadriannulatus | 18 | 7 | 2.57 | 22 | 35 | 0.629 | 0.008 | 0.334 |
| Gr18 | arabensis-quadriannulatus | 6 | 2 | 3.00 | 20 | 51 | 0.392 | 0.014 | 0.468 |
| Gr18 | coluzzii-quadriannulatus | 5 | 1 | 5.00 | 15 | 52 | 0.288 | 0.005 | 0.609 |
| Gr21 | gambiae-quadriannulatus | 2 | 0 | NA | 32 | 98 | 0.327 | 0.065 | 0.754 |
| Gr21 | merus-quadriannulatus | 7 | 8 | 0.875 | 12 | 49 | 0.245 | 0.045 | 0.270 |
| Gr26 | coluzzii-merus | 10 | 6 | 1.67 | 31 | 84 | 0.369 | 0.008 | 0.355 |
| Gr26 | gambiae-merus | 9 | 6 | 1.50 | 28 | 88 | 0.318 | 0.011 | 0.359 |
| Gr27 | coluzzii-merus | 15 | 5 | 3.00 | 56 | 58 | 0.966 | 0.050 | 0.259 |
| Gr35 | gambiae-melas | 9 | 15 | 0.6 | 21 | 98 | 0.214 | 0.051 | 0.199 |
| Gr35 | gambiae-merus | 5 | 8 | 0.625 | 17 | 106 | 0.160 | 0.037 | 0.246 |
| Gr47 | coluzzii-melas | 5 | 3 | 1.67 | 17 | 62 | 0.274 | 0.023 | 0.410 |
| Gr47 | gambiae-melas | 5 | 3 | 1.67 | 32 | 35 | 0.914 | 0.049 | 0.351 |
| Gr47 | melas-quadriannulatus | 7 | 7 | 1.00 | 12 | 42 | 0.286 | 0.051 | 0.278 |
| Gr48 | arabensis-quadriannulatus | 12 | 6 | 2.00 | 26 | 40 | 0.650 | 0.060 | 0.273 |
| Gr48 | gambiae-quadriannulatus | 3 | 0 | NA | 56 | 95 | 0.589 | 0.054 | 0.629 |
| Gr50 | arabensis-gambiae | 7 | 3 | 2.33 | 41 | 73 | 0.562 | 0.045 | 0.340 |
| Gr50 | gambiae-quadriannulatus | 7 | 4 | 1.75 | 42 | 88 | 0.477 | 0.049 | 0.313 |
| Gr51 | arabensis-merus | 19 | 7 | 2.71 | 33 | 36 | 0.917 | 0.037 | 0.253 |
| Gr51 | coluzzii-merus | 16 | 3 | 5.33 | 54 | 66 | 0.818 | 0.002 | 0.392 |
| Gr51 | merus-quadriannulatus | 11 | 4 | 2.75 | 30 | 49 | 0.612 | 0.021 | 0.354 |
| Gr56-RD | arabensis-merus | 10 | 5 | 2.00 | 17 | 38 | 0.447 | 0.017 | 0.358 |
| Gr56-RD | coluzzii-merus | 10 | 4 | 2.50 | 23 | 57 | 0.404 | 0.005 | 0.427 |
| Gr56-RD | gambiae-merus | 11 | 3 | 3.67 | 30 | 83 | 0.361 | 0.000 | 0.520 |
| Gr59 | gambiae-merus | 13 | 11 | 1.18 | 22 | 49 | 0.449 | 0.052 | 0.232 |
| Gr60 | coluzzii-melas | 13 | 7 | 1.86 | 16 | 34 | 0.471 | 0.016 | 0.330 |
| Gr60 | coluzzii-quadriannulatus | 8 | 4 | 2.00 | 22 | 42 | 0.524 | 0.053 | 0.323 |

In addition to the significant excess of fixed replacement substitutions identified when comparing *An. quadriannulatus* to *An. arabiensis* and *An. coluzzii*, *Gr18* also has a positive DoS (0.342) between *An. gambiae* s.s. and *An. quadriannulatus*, although the uncorrected p-value was not below 0.05 for this comparison. Interestingly, *Gr18* also shows evidence of recovery from a selective sweep in *An. gambiae* s.s. In addition, haplotype diversity (HD) for *Gr18* is lower in *An. coluzzii* than in *An. quadriannulatus*, although nucleotide diversity (π) is similar. Despite its known ligand, *Gr18* is lowly expressed in both *An. coluzzii* and *An. Quadriannulatus* chemosensory tissues *Gr48* has a marginally significant (p = 0.054) excess of fixed replacement substitutions between *An. gambiae* and *An. quadriannulatus*, a nonsignificant excess between *An. coluzzii* and *An. quadriannulatus*, and a marginally significant excess (p = 0.06) between *An. arabiensis* and *An. quadriannulatus*. *Gr48* has no known ligand but is relatively highly expressed in male *An. quadriannulatus* labella (unpublished data). Furthermore, π and HD are both lower in *An. quadriannulatus* than in *An. gambiae* s.s.

*Gr60* has a marginally significant (p = 0.053) excess of fixed replacement substitutions between *An. coluzzii* and *An. quadriannulatus* and a nonsignificant excess between *An. gambiae* and *An. quadriannulatus*, but a nonsignificant lack of fixed replacement substitutions between *An. arabiensis* and *An. quadriannulatus*. *Gr60* has no known ligand but is relatively highly expressed in male *An. quadriannulatus* maxillary palps[6]. Like *Gr48*, both π and HD are lower than in *An. coluzzii*.

Finally, *Gr4* has a significant excess of fixed replacement substitutions between *An. arabiensis* and *An. quadriannulatus*, but no fixed replacement substitutions between the latter and either *An. coluzzii* or *An. gambiae*. This gene is lowly expressed in both male and female *An. coluzzii* labella, as well as female *An. quadriannulatus* labella, but is highly expressed in male *An. quadriannulatus* labella (unpublished data).

### Purifying Selection

Ten *Grs* in the *An. gambiae* complex show evidence of purifying selection, as determined by the MK test identifying a significant lack of fixed replacement substitutions (**Table 4**). In addition, two other *Grs* show marginally significant evidence of purifying selection as determined by the MK test. Only one *Gr* meeting the criteria identified above is significant: *Gr19* shows evidence of purifying selection between *An. arabiensis* and *An. quadriannulatus*.

**Table 4.** Grs with significant (or near-significant lack of fixed substitutions and negative direction of selection (DoS), suggestive of purifying selection.

| <u>Gene</u> | <u>Species Pair</u> | <u>dN</u> | <u>dS</u> | <u>dN/dS</u> | <u>pN</u> | <u>pS</u> | <u>pN/pS</u> | <u>p-value</u> | <u>DoS</u> |
| --- | --- | --- | --- | --- | --- | --- | --- | --- | --- |
| Gr8 | *melas-merus* | 8 | 29 | 0.276 | 14 | 10 | 1.40 | 0.006 | -0.367 |
| Gr13-RA | *coluzzii-melas* | 1 | 7 | 0.143 | 32 | 27 | 1.19 | 0.054 | -0.417 |
| Gr19 | *arabiensis-quadriannulatus* | 0 | 7 | 0.000 | 28 | 46 | 0.609 | 0.050 | -0.378 |
| Gr22 | *melas-merus* | 1 | 17 | 0.059 | 7 | 11 | 0.636 | 0.041 | -0.333 |
| Gr33 | *melas-merus* | 0 | 19 | 0.000 | 3 | 9 | 0.333 | 0.049 | -0.250 |
| Gr37-RE | *arabiensis-quadriannulatus* | 1 | 9 | 0.111 | 21 | 21 | 1.00 | 0.032 | -0.400 |
| Gr38 | *arabiensis-merus* | 4 | 21 | 0.190 | 17 | 17 | 1.00 | 0.012 | -0.340 |
| Gr38 | *coluzzii-merus* | 2 | 17 | 0.118 | 25 | 28 | 0.893 | 0.005 | -0.366 |
| Gr38 | *gambiae-merus* | 2 | 16 | 0.125 | 29 | 38 | 0.763 | 0.013 | -0.322 |
| Gr39 | *arabiensis-quadriannulatus* | 0 | 14 | 0.000 | 16 | 25 | 0.640 | 0.005 | -0.390 |
| Gr39 | *coluzzii-quadriannulatus* | 0 | 10 | 0.000 | 25 | 39 | 0.641 | 0.013 | -0.391 |
| Gr39 | *gambiae-quadriannulatus* | 0 | 8 | 0.000 | 35 | 47 | 0.745 | 0.021 | -0.427 |
| Gr44-RE | *arabiensis-quadriannulatus* | 2 | 12 | 0.167 | 56 | 52 | 1.08 | 0.010 | -0.376 |
| Gr44-RE | *coluzzii-quadriannulatus* | 2 | 14 | 0.143 | 36 | 45 | 0.800 | 0.023 | -0.319 |
| Gr44-RE | *gambiae-quadriannulatus* | 2 | 12 | 0.167 | 56 | 66 | 0.848 | 0.025 | -0.316 |
| Gr45 | *coluzzii-quadriannulatus* | 0 | 4 | 0.000 | 34 | 28 | 1.21 | 0.050 | -0.548 |
| Gr54 | *arabiensis-merus* | 11 | 22 | 0.500 | 23 | 13 | 1.77 | 0.016 | -0.306 |
| Gr60 | *melas-merus* | 16 | 22 | 0.727 | 15 | 6 | 2.50 | 0.055 | -0.293 |

### DH Test

Selective sweeps can be detected by significantly negative Tajima’s D values, as well as significantly negative Fay and Wu’s H values. However, since both tests are subject to biases from demographic forces, we only consider genes with significantly negative values on the DH test, which is a combination of the two tests and was designed to be robust to the influence of demographic factors [118], to show strong evidence of sweep. As the DH test was only run on fully genotyped sequences, far fewer sequences were available for analysis than for the MK test. For *An. coluzzii* and *An. gambiae* s.s., data were available for most *Grs*: 50 and 53, respectively. *An. quadriannulatus* data was available for 36 *Grs*, while much less data was available for the other species: three *Grs* in *An. arabiensis*, seven in *An. melas*, and six in *An. merus*. As above, *An. melas* and *An. merus* are included in data tables but not discussed in the text. Results of all tests are shown in **Supplemental Tables 3 and 4**.

A total of five *Grs* show significant evidence of a potential selective sweep based on the DH test, while an additional three *Grs* show nearly significant (0.05 ≥ p ≥ 0.06) evidence thereof (**Table 5**). In *An. coluzzii*, the highly expressed sugar receptor, *Gr17*, is the only *Gr* showing significant evidence of sweep. The TCS network of *Gr17* in *An. coluzzii* is somewhat star-shaped, but there is no central high-frequency haplotype, which could be a consequence of the length of time following the sweep (**Figure 3**). In *An. quadriannulatus*, the lowly expressed sugar receptor *Gr19* shows significant evidence of a potential selective sweep. The TCS network features a clear star shape with central high-frequency haplotype surrounded by lower-frequency haplotypes, which is a hallmark of sweep (**Figure 4**).

**Figure 3.**
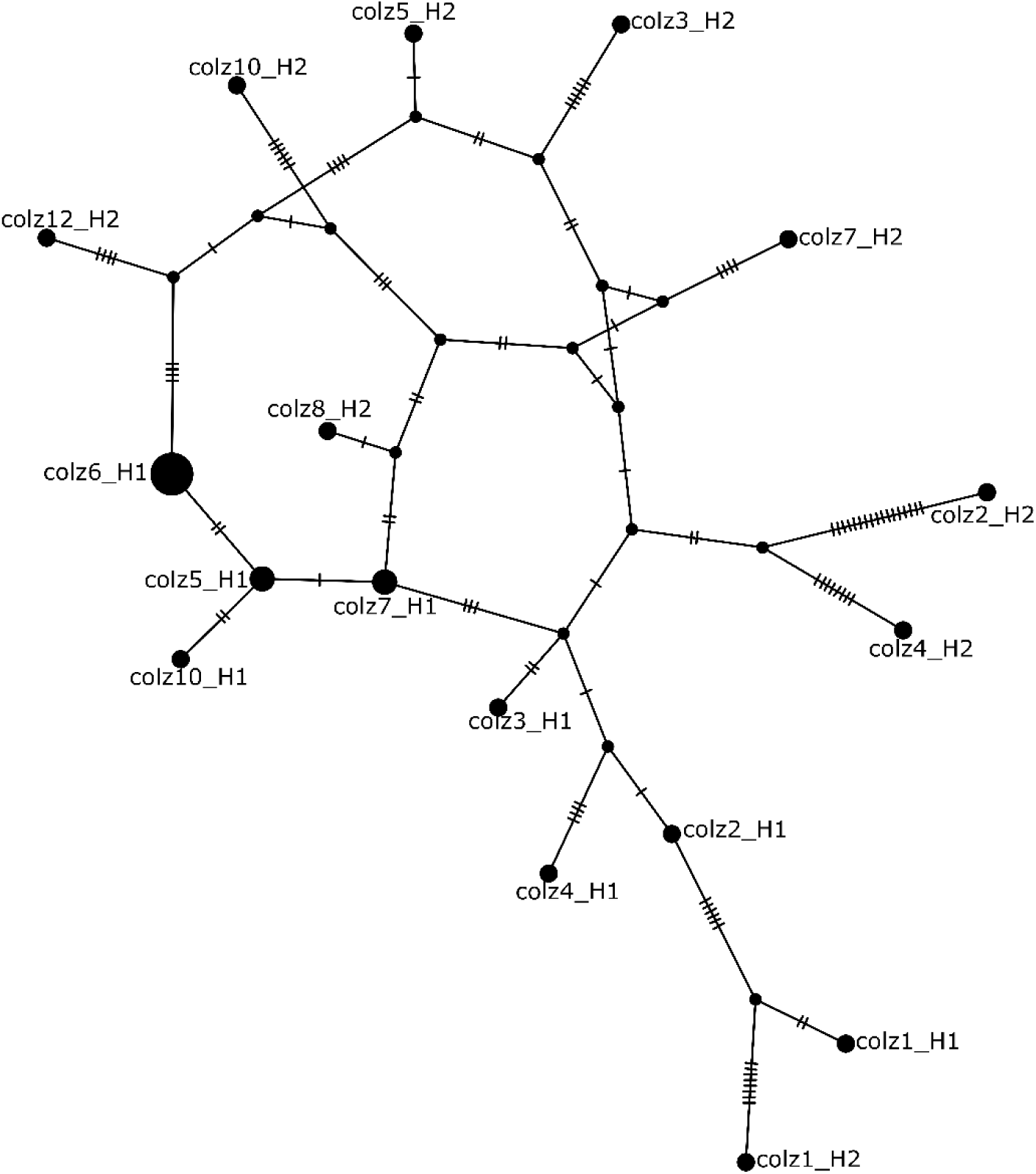
TCS Haplotype Network of the sugar receptor Gr17 in An. coluzzii. Each hashed line represents one nucleotide substitution and haplotype nodes are weighted by frequency. There are 17 haplotypes among 24 sequences (π = 0.008 and HD = 0.938).

**Figure 4.**
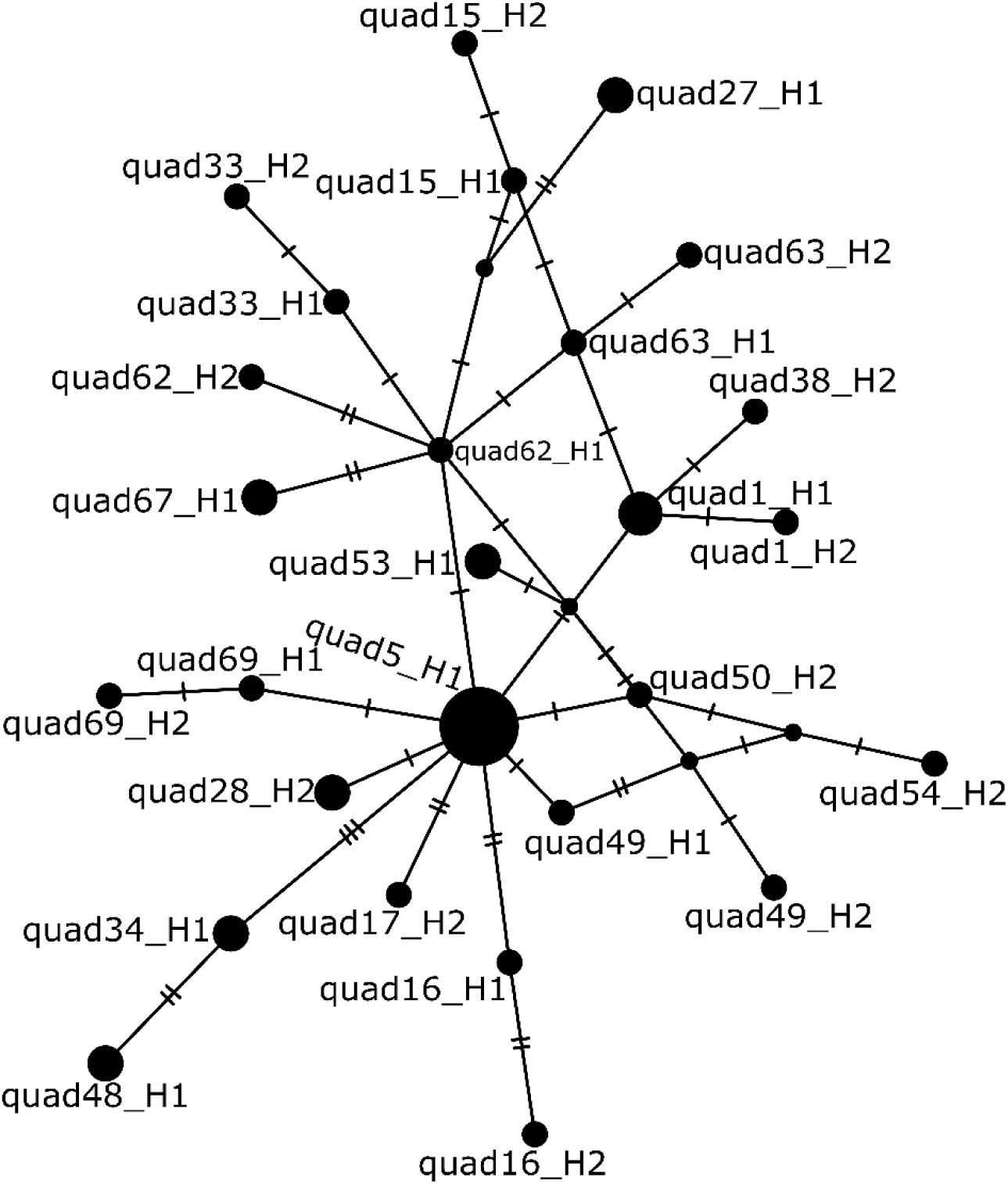
TCS Haplotype Network of the sugar receptor Gr19 in An. quadriannulatus. Each hashed line represents one nucleotide substitution and haplotype nodes are weighted by frequency. There are 16 haplotypes among 44 sequences (π = 0.002 and HD = 0.943).

**Table 5.** Grs with significant (or near significant) DH values, suggestive of a selective sweep.

| <u>Gene</u> | <u>Species</u> | <u>DH</u> | <u>$\pi$</u> | <u>Haplotype Diversity (HD)</u> |
| --- | --- | --- | --- | --- |
| *Gr3* | *An. melas* | 0.044 | 0.001 | 0.632 |
| *Gr11* | *An. melas* | 0.046 | 0.001 | 0.556 |
| *Gr12* | *An. melas* | 0.052 | 0.002 | 0.667 |
| *Gr17* | *An. coluzzii* | 0.047 | 0.008 | 0.938 |
| *Gr19* | *An. quadriannulatus* | 0.007 | 0.002 | 0.943 |
| *Gr36* | *An. gambiae* | 0.056 | 0.008 | 0.998 |
| *Gr41* | *An. quadriannulatus* | 0.019 | 0.002 | 0.94 |
| *Gr59* | *An. gambiae* | 0.054 | 0.008 | 0.988 |

### E Test (Recovery from Selective Sweep or Background Selection)

Two *Grs* in *An. gambiae* s.s. show significant E test values, which detect an excess of low-frequency polymorphisms suggestive of recovery from a selective sweep or background selection: *Gr18*, and *Gr24* (**Table 6**). *Gr18* encodes a lowly-expressed sugar receptor, but is adjacent to *Gr17*, which is a very highly expressed sugar receptor. In addition to its excess of low-frequency variants in *An. gambiae* (**Supplemental Figure X)**, it has an excess of fixed replacement substitutions between *An. quadriannulatus* and both of the other major vectors, *An. arabiensis* and *An. coluzzii*. The CO_2_ receptor *Gr24* also has a significant excess of low-frequent variants (**Supplemental Figure X**), which is surprising due to the high sequence conservation of the CO_2_ receptors and the importance of CO_2_ as a host cue.

**Table 6.**
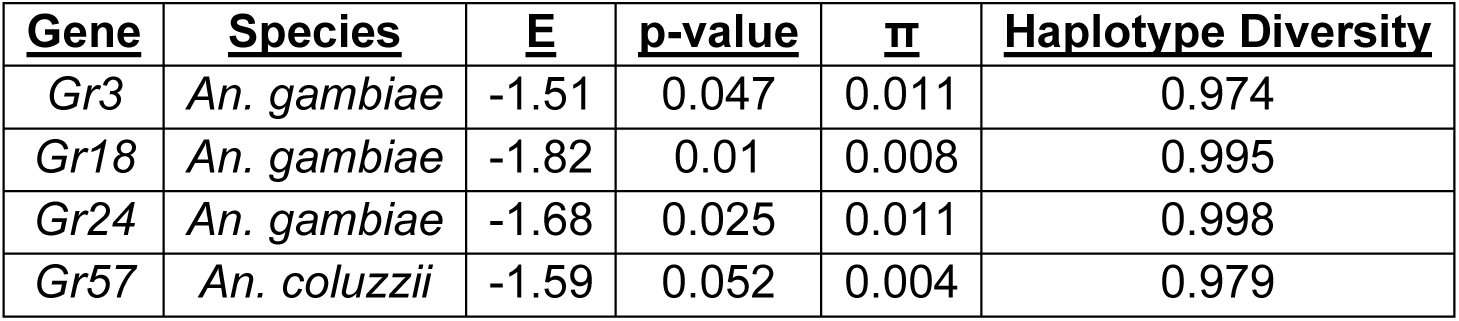
Grs with significant (or near significant) E-test values, indicative of recovery from a selective sweep.

| <u>Gene</u> | <u>Species</u> | <u>E</u> | <u>p-value</u> | <u>$\pi$</u> | <u>Haplotype Diversity</u> |
| --- | --- | --- | --- | --- | --- |
| *Gr3* | *An. gambiae* | -1.51 | 0.047 | 0.011 | 0.974 |
| *Gr18* | *An. gambiae* | -1.82 | 0.01 | 0.008 | 0.995 |
| *Gr24* | *An. gambiae* | -1.68 | 0.025 | 0.011 | 0.998 |
| *Gr57* | *An. coluzzii* | -1.59 | 0.052 | 0.004 | 0.979 |

## Discussion

In this study, we examined the selective forces acting on *Grs* in the *An. gambiae* complex to determine whether they are likely to play a role in differing vertebrate host preference between constituent species. Our analyses suggest that sixteen *Grs*, six of which either have known functions or are highly expressed in chemosensory organs, are primarily under positive selection based on comparisons between *An. arabiensis*, *An. coluzzii*, *An. gambiae* s.s., *An. melas*, *An. merus*, and *An. quadriannulatus* using the McDonald-Kreitman test. Our analyses further suggest purifying selection in twelve *Grs*. Furthermore, the DH test indicates that eight *Gr*s are under the influence of sweep and the E test indicates that four *Gr*s are either recovering from either sweep or subject to background selection. While the evolution of *Drosophila Grs* and its relationship to ecological adaptations has been extensively studied [57, 63–66], and there have been several studies on the evolution of *Grs* in Lepidoptera [67, 69, 70, 75, 135], the mosquito literature on the evolution of chemosensory genes is much more limited [77, 78]. As such, we can compare what we know of *Gr* evolution in other lineages while incorporating the evolutionary and ecological contexts of the six *An. gambiae* s.l. species. There is evidence that *Grs* are under both positive and purifying selection in the *An. gambiae* complex, which may have contributed to the development of species-specific behavioral ecology, as is seen in other taxa, most notably the *An. farauti* complex [78]. As in other taxa [57, 64, 74, 75], in most species comparisons within the *An. gambiae* complex, most *Gr*s have negative DoS values, consistent with purifying selection.

Selection tests, such as branch tests, that can test for selection on specific lineages within a phylogeny are available [136]. These tests examine if an excess of replacement substitutions is found along specific, pre-defined lineages of interest, of even if specific sites within a lineage have such an excess. We applied the branch test and branch-site test to the chemosensory genes within the *An. gambiae* complex, but found that the signal was biased by the presence of ancestral polymorphisms, and that no reliable inference could be made. Therefore, these tests are not included in the present paper.

In those *Grs* that show signatures of positive selection, biological significance is mostly unclear, as they have unknown ligands, excluding two sugar receptors (*Grs* 18 and 21). However, homologs of three others (*Gr4*, *Gr59*, *Gr60*) were also identified as candidates for being correlated with host preference in the *An. farauti* complex [78]. Both sugar receptors are expressed in both male and female labella [8] (unpublished data). Two of the *Grs* showing evidence of positive selection between *An. quadriannulatus* and *An. coluzzii* (*Gr60*) and *An. gambiae* s.s. (*Gr48*) are male-biased and *An. quadriannulatus*-biased in the maxillary palps [6] and labella (unpublished data), respectively. While females engage in several sex-specific behaviors such as blood-feeding, host-seeking, and oviposition, the only male-specific behaviors are swarming [137] and mating. The antennal fibrillae are known to play an important role in male detection of auditory cues in both swarming and close-range mating behaviors in the *An. gambiae* complex, and the male claspers recognize if females have mated, but other organs have not been implicated in male mating [137, 138]. While neither the maxillary palps nor labella have been established as playing a role in mating biology in *Anopheles*, there is evidence that the maxillary palp detects female inhibitory cues in *Drosophila* [139], which raises the possibility that a similar phenomenon might occur in *Anopheles*, particularly as *Drosophila Grs* expressed in the labellum and tarsi are known to play a role in inhibiting male-male courtship [140]. The labellum is well-established in a male mating role in *Drosophila* [141]. While work on the role (if any) of the tarsi and male mouthparts in anopheline mating is ongoing, there is evidence that females in mixed-species swarms mate assortatively [142], although males are not believed to do so [138], despite the ability to detect females who have already mated.

Aside from *Grs* 18, 21, 48, and 60, most other *Grs* with MK test results consistent with positive selection are lowly expressed in chemosensory organs or show evidence of selection in species with uncharacterized transcriptomes. As such, it is more difficult to explain their biological relevance, but they may be expressed in other organs [40, 43–51]. With the exception of the CO_2_ and sugar receptors, the ligands of anopheline *Gr*s are unknown. This is in contrast to *Drosophila*, where they are known to also perceive bitter compounds and cuticular hydrocarbons, as well as mediate light and heat avoidance [31, 32, 37–42, 50, 51].

### Furthermore, *Grs* are difficult to deorphanize [52], and most anopheline *Grs* do not have known *Drosophila* homologs

With respect to the *Gr*s that show signatures of purifying selection, one (*Gr19*) is a sugar receptor, one (*Gr22*) is a CO_2_ receptor, and homologs of three are candidates for a host preference association in the *An. farauti* complex (*Gr13*, *Gr22*, and *Gr39*). Since only *Gr19* and *Gr22* have known ligands, it is not currently possible to explain why deleterious mutations would be purged in the other *Gr*s under purifying selection. It is intuitive that deleterious mutations in *Gr22* would result in a fitness cost, but less clear why this was only detected in the comparison between *An. melas* and *An. merus*.

Eight *Grs* have DH test results consistent with recent selective sweeps (**Table 5**). As discussed above, for *Grs* that are expressed in the mouthparts of either *An. coluzzii* or *An. quadriannulatus*, this could mean that these *Gr*s are involved in the adaptation to hosts. However, of these eight Grs, only the sugar receptor *Gr17* is highly expressed. Its expression, like that of *Gr18*, is not sex-biased, so the adaptation of these genes presumably does not underly any sex-biased behaviors. An obvious role for *Gr17* is nectar-feeding. Nectar is the only resource fed on by adult males but also prolongs the lives and reduces blood-feeding frequency of adult females [143]. While the other *Grs* with significant DH test results consistent with selective sweeps are lowly expressed, many of them are located near genes encoding critical cellular functions or other *Grs* which are more highly expressed and also show evidence of positive selection. These selective signatures could therefore be due to hitchhiking. However, interestingly, four of these *Gr*s (*Gr12*, *Gr36*, *Gr41*, *Gr59*) have homologs that are candidates for an association with vertebrate host preference in the *An. farauti* complex, with the strongest evidence for such an association in *Gr36* (based on phylogenetic analysis) and *Gr41* (which shows evidence of intensified selection in all zoophilic lineages within this complex) [78]. Since *Gr41* shows significant evidence of sweep in *An. quadriannulatus*, it is possible that this *Gr* plays a role in zoophily in both species complexes.

Four *Grs* show E test results consistent with recovery from a selective sweep, although the E test is also sensitive to background selection [118]. *Gr57*, the *Gr* that may play a role in anthropophily in both the *An. farauti* and *An. gambiae* complexes [78], has a near-significant result in *An. coluzzii*. Besides *Gr18*, the CO_2_ receptor *Gr24* has a significant E test value in *An gambiae* s.s. The CO_2_ receptors are highly conserved across insects [144], and there are no fixed differences in *Gr24* between *An. arabiensis*, *An. coluzzii*, *An. gambiae* s.s., and *An. quadriannulatus*, although there are some when compared to *An. melas* and *An. merus*. CO_2_ is thought to matter less as a host-seeking cue in the anthropophilic members of the complex than in more zoophilic or generalist species in the complex [87]. This lack of fixed differences suggests background selection against deleterious mutations as an explanation for this significant E test result.

While we have high-quality variant data from *An. coluzzii* and *An. gambiae* s.s. for these 57 *Grs*, data for the other four species are of a much lower quality, as indicated by the overall sequencing coverage, the number of variants detected, and the number of complete sequences. While we have sufficient data to draw conclusions about evolutionary patterns, particularly with respect to the zoophilic non-vector *An. quadriannulatus*, a more comprehensive analysis of evolutionary patterns that differentiate the other species in the complex from one another would require either deeper coverage of the whole genome sequences of the remaining species, or a more targeted sequencing approach, such as amplicon sequencing or molecular inversion probes, which would allow precise sequencing of the *Grs* at a much greater depth.

Of the *Gr*s showing significant signatures of selection, *Gr41*, *Gr57*, *Gr59*, and *Gr60* are the most attractive targets for further study, given that they have previously been identified as candidates for association with vertebrate host preference in *An. farauti* s.l. and significantly differ from neutral expectations in this analysis when comparing anthropophilic to zoophilic species (*Gr60*), within anthropophilic species (*Gr57*, *Gr59*), or within zoophilic species (*Gr41*). *Gr59* is expressed in both male and female antennae and labella of both *An. coluzzii* and *An. quadriannulatus* [6, 8, 9, 12], so could be a viable knockdown target for behavioral or electrophysiological assays to better elucidate its biological role. Similarly, *Gr60* is *An. quadriannulatus*-biased in both male and female maxillary palps [6, 9], and could be treated in the same way. Neither *Gr41* nor *Gr57* have been detected in *An. coluzzii* or *An. quadriannulatus* mouthparts, so determining their functional significance will first require studies to determine if they are expressed in other organs or life stages before any more targeted work can occur.

The sugar receptors *Grs* 14-21 are closely clustered on the 2R chromosome, and three of them show evidence of directional selection in either an anthropophilic species or *An. quadriannulatus*. Though their function is not understood to the same degree as in *Drosophila*, the potential presence of positive selection in them merits further study, as these may be involved in the recent adaptive divergence between species in this complex, including potentially divergence in host preference or preferred habitat, as it could relate to the ability to successfully exploit preferred nectar resources, or to evaluate sugar sources which are found in greater abundance near preferred vertebrate hosts.

There are several limitations to our current knowledge that preclude more specific hypotheses for the biological importance of most of the *Gr*s that show significant signatures of selection in this study. First, transcriptomic data is lacking for species other than *An. coluzzii* and *An. quadriannulatus*, as well as for other organs that are known to express *Gr*s in *Drosphila*, including the tarsi, wings, and internal organs such as the brain and midgut [40, 43–51]. There is a similar dearth of transcriptomic data on *Gr*s in larval *Anopheles*, though studies have characterized their *Or* and *Ir* repertoires, as well as behavioral responses to odorants [145–147]. Research on *Aedes aegypti* larvae has shown that they rely on chemokinesis to navigate chemical gradients, and has further suggested that they likely rely on *Grs* and *Irs* rather than *Ors* to do so [148]. As such, characterization of *Gr* expression profiles in larval *Anopheles* could potentially illuminate biological meaning for some lowly expressed *Grs* in adults.

Even within *Gr*s that have known ligands or expression profiles, signatures of selection do not necessarily reflect selection on the *Gr* in question, and may instead reflect hitchhiking due to other linked genes that are unrelated to vertebrate host preference. Therefore, functional studies of these *Gr*s are needed to determine what role, if any, they play in determining host preference. In addition, GRs are frequently co-expressed [22, 149–151], meaning that it is difficult to disentangle their roles from one another.

## Conclusions

In this study, we have presented the first analysis of the molecular evolution of gustatory receptors within the *Anopheles gambiae* complex of malaria vectors, including six species with genomic data available. We have identified 16 *Grs* that with signatures suggesting positive selection within this complex, six of which either have known functions or are highly expressed in the chemosensory organs of either the highly anthropophilic *An. coluzzii* or the zoophilic *An. quadriannulatus*. In addition, we have identified twelve *Grs* that may be undergoing purifying selection and twelve *Grs* that may be under the influence of sweep. Based on signatures of selection and gene expression data, this study identifies four *Grs* as possible candidates in the adaptation to distinct vertebrate hosts in the *An. gambiae* complex, as may have occurred with homologous genes in the *An. farauti* complex. However, further elucidating this question will require additional study.

## Declarations

Ethics approval and consent to participate: Not applicable.

## Consent for publication

Not applicable.

## Availability of data and materials

All sequences used in this study are publicly available from NCBI. A full list of accession numbers can be found in **Supplementary Table 5**.

## Authors’ contributions

The project was conceived by MAS. ZRPH and MAS designed the study, analyzed the data, and wrote the final manuscript. Bioinformatic pipelines were developed by ZRPH. All authors read and approved the final manuscript.

## Acknowledgements

Giridhar Athrey provided helpful advice on the variant calling pipeline. Adrian Castellanos, Curtis Cooper, and Jenna Hull helped diagnose coding errors. This work was conducted as part of ZRPH’s doctoral dissertation, which was supervised by a committee including Giridhar Athrey, J. Spencer Johnston, Christine Merlin, and Aaron M. Tarone.

